# Advancing Clinical Response Against Glioblastoma: Evaluating SHP1705 CRY2 Activator Efficacy in Preclinical Models and Safety in Phase I Trials

**DOI:** 10.1101/2024.09.17.613520

**Authors:** Priscilla Chan, Yoshiko Nagai, Qiulian Wu, Anahit Hovsepyan, Seda Mkhitaryan, Jiarui Wang, Gevorg Karapetyan, Theodore Kamenecka, Laura A. Solt, Jamie Cope, Rex A. Moats, Tsuyoshi Hirota, Jeremy N. Rich, Steve A. Kay

## Abstract

**Background:** It has been reported that circadian clock components, Brain and Muscle ARNT-Like 1 (BMAL1) and Circadian Locomotor Output Cycles Kaput (CLOCK), are uniquely essential for glioblastoma (GBM) stem cell (GSC) biology and survival. Consequently, we developed a novel Cryptochrome (CRY) activator SHP1705, which inhibits BMAL1-CLOCK transcriptional activity.

**Methods:** We analyzed buffy coats isolated from Phase 1 clinical trial subjects’ blood to assess any changes to circadian, housekeeping, and blood transcriptome-based biomarkers following SHP1705 treatment. We utilized GlioVis to determine which circadian genes are differentially expressed in non-tumor versus GBM tissues. We employed *in vitro* and *in vivo* methods to test the efficacy of SHP1705 against patient-derived GSCs and xenografts in comparison to earlier CRY activator scaffolds. Additionally, we applied a novel-REV-ERB agonist SR29065, which inhibits *BMAL1* transcription, to determine whether targeting both negative limbs of the circadian transcription-translation feedback loop (TTFL) would yield synergistic effects against various GBM cells.

**Results:** SHP1705 is safe and well-tolerated in Phase I clinical trials. SHP1705 has increased selectivity for the CRY2 isoform and potency against GSC viability compared to previously published CRY activators. SHP1705 prolonged survival in mice bearing GBM tumors established with GSCs. When combined with the novel REV-ERB agonist SR29065, SHP1705 displayed synergy against multiple GSC lines and differentiated GSCs (DGCs).

**Conclusions:** These demonstrate the efficacy of SHP1705 against GSCs, which pose for GBM patient outcomes. They highlight the potential of novel circadian clock compounds in targeting GBM as single agents or in combination with each other or current standard-of-care.

**KEY POINTS:**

- SHP1705 is a novel CRY2 activator that has shown success in Phase 1 safety trials
- SHP1705 has a significantly improved efficacy against GSCs and GBM PDX tumors
- Novel REV-ERB agonist SR29065 and SHP1705 display synergistic effects against GSCs

**IMPORTANCE OF THE STUDY:** *CRY2* is decreased in GBM tissues compared to *CRY1* suggesting that promoting CRY2 activity will be an efficacious GBM treatment paradigm. SHP1705, a CRY2 activator that has shown success in Phase 1 safety trials, has significantly improved preclinical efficacy. Novel REV-ERB agonist SR29065 displays synergistic effects against diverse GBM cells.

## INTRODUCTION

GBM is the most common primary brain tumor type with a poor prognosis and limited effective treatment options especially after recurrence.^1^ GBM has a median survival time of 15 months with the current standard-of-care or the Stupp protocol.^2^ This consists of an aggressive regimen of maximal surgical resection followed by concurrent temozolomide (TMZ) chemotherapy and radiation then adjuvant TMZ.^3^ One of the major challenges in treating GBM is the presence of GBM stem cells (GSCs), which have tumor-initiating properties, secrete angiogenic factors, induce an immune suppressive environment, are highly invasive, and are resistant to both chemotherapy and radiation. GSCs can remain in and well beyond the margins of the surgical cavity following tumor resection and drive tumor recurrence.^4–6^

Core circadian clock components BMAL1 and CLOCK are essential to GSC biology. BMAL1 and CLOCK chromatin binding sites were shown to be increased compared to neural stem cells (NSCs), allowing for BMAL1-CLOCK-driven expression of stemness and metabolic genes in GSCs.^7^ BMAL1 and CLOCK drive expression of the chemokine Olfactomedin-Like 3 (OLFML3) in GSCs to recruit immune suppressive microglia and drive angiogenesis.^8,9^ Perturbation of either BMAL1 or CLOCK genetically or pharmacologically resulted in specific cell death in GSCs that was not seen in differentiated GSCs (DGCs) or noncancerous cells; downregulated stemness markers, the Citric Acid (TCA) cycle genes, and OLFML3; and decreased tumor volume and increased survival in GBM patient derived xenograft (PDX) mouse models established with GSCs.^7–9^

Circadian rhythms in mammals are generated by cycling in gene expression that is driven by BMAL1 and CLOCK transcriptional output in individual cells under tight control of a TTFL. This results in roughly a 24hr period in behavioral and physiological output. BMAL1 and CLOCK form a heterodimer (BMAL1-CLOCK) and converge upon the E-box motifs of clock-controlled gene (CCG) promoters to drive cyclical gene expression. BMAL1-CLOCK also control the expression of their positive and negative regulators. One TTFL loop of the clock consists of Cryptochrome 1/2 (CRY1/2) and Period 1/2 (PER1/2), which form a heterodimer that inhibits the BMAL1-CLOCK transcriptional loop. In another TTFL loop, REV-ERBαβ inhibits while Retinoic Acid-Related Orphan Receptor α/β/γ (RORα/β/γ) promotes the transcription of *BMAL1* by acting upon the ROR-element (RORE) in its promotor **(Fig. 1A)**. This TTFL network and their post-translational modifications generate rhythmicity in key bodily functions.^10,11^

**Figure 1.**
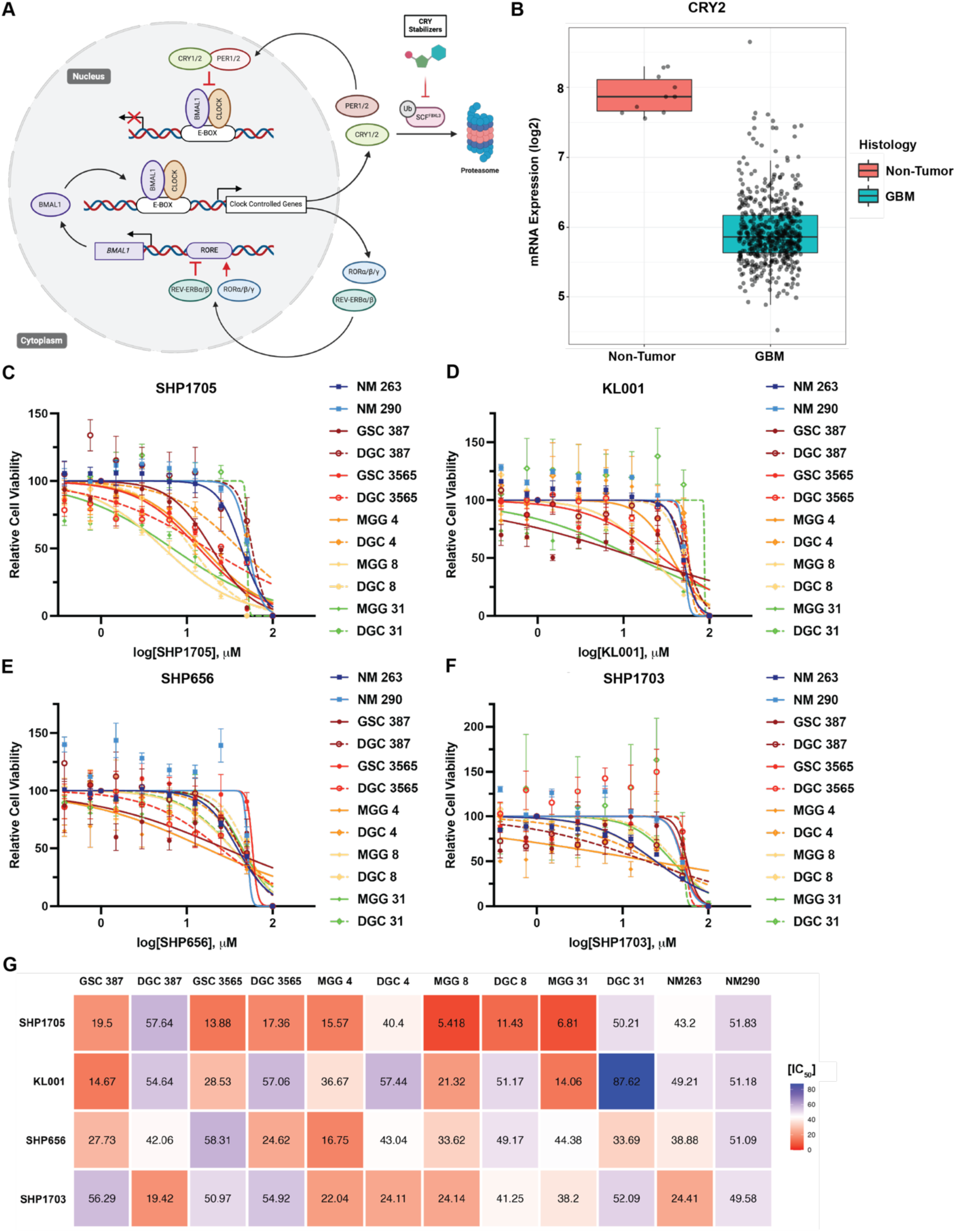
SHP1705 displays significantly improved anti-GSC effects compared to earlier CRY activator scaffolds. **(A)** Mechanism of the mammalian clock and CRY activators. BMAL1 and CLOCK form a heterodimer and converge upon the E-box motifs of clock-controlled gene promoters to drive their transcription. BMAL1 and CLOCK also drive the expression of their positive and negative regulators. CRYs and PERs form a heterodimer and inhibit the transcriptional activity of BMAL1-CLOCK. RORα/β/γ promote while REV-ERBαβ repress the expression of *BMAL1* by acting upon the RORE of its promoter. CRY activators prevent FBXL3-mediated ubiquitination of CRY1/2 proteins and their downstream degradation. Consequently, CRY1/2 can continue to form a heterodimer with PER1/2 and repress BMAL1-CLOCK transcriptional activity. **(B)** *CRY2* mRNA expression of non-tumor and GBM tissues from adult TCGA_GBM HG-U133A data plotted and analyzed with GlioVis. **(C-G)** Cell viability analysis of NM cells (solid blue lines), GSCs (solid lines), or DGCs (dotted lines of matched GSC solid lines) following **(C)** SHP1705, **(D)** KL001, **(E)** SHP656, or **(F)** SHP1703 treatment for 3 days (*n* = 3-4 biologically independent samples). **(G)** Heatmap of summarized IC_50_ values following CRY activator treatment for 3 days in control NM cells, GSCs, and DGCs.

BMAL1 and CLOCK are transcription factors that, to date, have been difficult to directly target pharmacologically. However, small molecules have been developed that indirectly modulate their function or transcription by directly engaging with their negative regulators CRY and REV-ERB.^11–13^ One such type of circadian clock compounds that have been developed by our group are the CRY activators. These were identified through a series of high throughput luciferase-based reporter assays that monitored changes in *Bmal1* (*Bmal1-dLuc*) and *Per2* (*Per2-dLuc*) gene expression cycling. CRY activators compete with F-Box and Leucine-Rich Repeat Protein (FBXL3) in the Flavin Adenine Dinucleotide (FAD) binding pockets of CRY1/2, therefore preventing ubiquitination and downstream proteasome degradation **(Fig. 1A)**. KL001 was the first reported CRY activator and, since then, bioavailable CRY isoform-selective scaffolds, such as SHP656 and SHP1703, have since been developed. We have previously shown that these compounds have anti-GSC effects in both *in vitro* and *in vivo* assays, and SHP656 and SHP1703, in particular, are selective for CRY2.^7,14–17^

CRY2 was found to be dysregulated in a glioma rat model and sensitize cells to apoptosis following radiation.^18^ These studies suggest that pharmacologically perpetuating CRY2 function can have favorable anti-cancer effects against GSCs and GBM tumors. Consequently, in this study we focused specifically on investigating the efficacy of a novel CRY2 activator scaffold, SHP1705.

## MATERIALS AND METHODS

### Derivation of Glioblastoma Stem Cells

GSC 387, GSC 3565, MGG 4, MGG 8, and MGG 31 were derived from patients as previously described.^35,36^ hGBM18 FMC MP1 generated from MGG 18 by adding in firefly luciferase and mCherry reporters and passaging the cells in mice to increase the growth rate for *in vivo* experiments.

### Derivation of Nonmalignant Neuronal Cultures

NM 263 and NM 290 cells were derived from surgical specimens from patients with epilepsy.^35^

### Cell Culture

GSC 387, GSC 3565, MGG 4, MGG 8, MGG 31, and hGBM18 FMC MP1 were cultured in Neurobasal-A medium without phenol (Gibco, Cat. #12349-015), B27 Supplement without Vitamin-A (Life Technologies, Cat. #12587-010), 1% penicillin/streptomycin (10,000 U/mL, Gibco, Cat. #15140-122), 1% GlutaMAX^TM^ (100X, Gibco, Cat. #35050-61), 1% Sodium Pyruvate (100 mM, Gibco, Cat. #11360-070), 0.02 ug/mL EGF (0.5 mg/mL dissolved in PBS, R&D, Cat. #236-EG-01M), and 0.02 ug/mL FGF (0.5 mg/mL dissolved in PBS, Cat. #4114-TC-01M) (complete Neurobasal-A media). NM cells were cultured in 1:1 ratio of Dubecco’s Modified Eagle Medium (DMEM) (Gibco, Cat. #11995-065), 10% fetal bovine serum (FBS, R&D Systems, Cat. #S11150), and 1% penicillin/streptomycin (10,000 U/mL, Gibco, Cat. #15140-122) and complete Neurobasal-A media. DGCs were differentiated from GSCs and maintained by adding 10% FBS (R&D Systems, Cat. #S11150) to complete Neurobasal-A media for at least 48 hrs. Cells were all incubated at 37°C and 5% CO_2_ and kept at low number of passages.

### Cell Viability Assays

Cells were plated in 96-well black, clear, flat bottom plates (Falcon, Cat. #353219) at 1,000 cells/100 μL/well. SHP1705 or SR29065 was dissolved in DMSO (Sigma-Aldrich, Cat. #D2650-100ML) to make a stock concentration of 10 mM. SHP1705 and/or SR29065 was added at the desired concentrations accordingly to the plate. Cells were incubated at 37°C and 5% CO_2_ for 3-4 days with clock compound. 50 μL of Cell-Titer Glo (Promega, Cat. # G7573) was added to each well at the time of reading, and the plate was incubated at room temperature and protected from light for 15 mins. Luminescence was read as a function of ATP levels by the Tecan Infinite Pro M200 plate reader. Values were plotted in GraphPad Prism to generate cell viability curves and determine IC_50_ values.

### qPCR

Cells were plated in a 6-well plate at 1×10^6^ cells/well with the appropriate cell medium and the indicated concentrations of SHP1705 for 24 hrs. The cell lysates were then collected and snap frozen. RNA was extracted with the Qiagen RNA extraction kit as instructed with the manufacture’s guidelines. RNA was reversed transcribed to cDNA with the iScript^TM^ cDNA Synthesis Kit (Bio-Rad, Cat. #170-8891). RT-qPCR was performed with the Bio-Rad CFX Opus 384 Real-Time PCR System using the SsoAdvanced Universal SYBR^®^ Green Supermix (Bio-Rad, Cat. #1725274) Cycle conditions as follows: 95°C (30 sec), 95°C (10 sec), 60°C (60 sec), repeat Steps 2-3 39X, 65°C (5 sec), and increase to 95°C (0.5°C/cycle). Primers were purchased from Integrated DNA Technologies and sequences (5’ → 3’) are listed below.

*18S rRNA* Forward: GCTTAAATTTGACTCAACACGGGA

*18S rRNA* Reverse: AGCTATCAATCTGTCAATCCTGTC

*PER2* Forward: TACGCTGGCCACCTTGAAGTA

*PER2* Reverse: CACATCGTGAGGCGCCAGGA

### U2OS Circadian Assays

U2OS *Bmal1-dLuc* and *Per2-dLuc* cells were generated as previously published.^37,38^ Cells were maintained in DMEM (Gibco, Cat. #11995-065), 10% FBS (R&D Systems, Cat. #S11150), and 1% penicillin/streptomycin (10,000 U/mL, Gibco, Cat. #15140-122) at 37°C and 5% CO_2_. Cells were plated at 10,000 cells/100 μL/well in a 96-well white, flat bottom plate (Corning, Cat. #3917) and incubated for 24 hrs. Media was changed to an explant media (DMEM (2X, Gibco, Cat. #12800-017), 1% penicillin/streptomycin (10,000 U/mL, Gibco, Cat. #15140-122), 1% GlutaMAX^TM^ (100X, Gibco, Cat. #35050-61), 1 mM D-Luciferin, potassium salt (PerkinElmer, Cat. #122799), 400 μM NaOH (100 mM, Fluka Cat. #72079-100ML), 10% FBS (R&D Systems, Cat. #S11150), and Cell Culture Grade Water (Corning, Cat. #25-055-CM). 2X DMEM was made mixing DMEM powder (Gibco, Cat. #12800-017), 10 mL HEPES (1 M, Gibco, Cat. #15630-080), 5 mL sodium bicarbonate (7.5%, Gibco, Cat. #25080-094), and 485 mL Cell Culture Grade Water (Corning, Cat. #25-055-CM) and sterilizing it with a vacuum filtration cup. SHP1705 or SR29065 was dissolved in DMSO (Sigma-Aldrich, Cat. #D2650-100ML) to make a stock concentration of 10 mM. SHP1705 and/or SR29065 was added at the desired concentrations to the explant media accordingly to the plate. Plates were covered with a clear plastic film and maintained at 37°C in the Tecan Infinite Pro M200 plate reader, where luminescence was read every 2 hrs for 120 hrs. Values were plotted in Excel to generate graphs. Amplitude, period, and phase were calculated with MultiCycle (Actimetrics) and two-way ANOVA was performed for statistical analysis.

### *Per2::Luc* Repression Assay

Wild type, *Cry1/Cry2* double knockout, *Cry1* knockout, and *Cry2* knockout fibroblasts harboring a *Per2::Luc* knock-in reporter^39^ were plated on a white, solid-bottom 384-well plate and cultured for 2 days to reach confluency, followed by the application of 500 nL of compounds (final 0.7% DMSO). After 2 days, the medium was replaced with BrightGlo (Promega, Cat. #E2620), and luminescence was recorded in Cytation3 plate reader (BioTek).

### *Per2::Luc* Cell Circadian Assay

Wild type, *Cry1/Cry2* double knockout, *Cry1* knockout, and *Cry2* knockout fibroblasts harboring a *Per2::Luc* knock-in reporter were plated on a white, solid-bottom 96-well plate and cultured for 2 days to reach confluency. The cells were treated with forskolin (final 10 µM) for 2 hrs. Then, the medium was replaced with explant medium [DMEM (Gibco, Cat. #12800-017) supplemented with 2% B27 (Gibco), 10 mM HEPES, 0.38 mg/mL sodium bicarbonate, 0.29 mg/mL L-glutamine, 100 units/mL penicillin, 100 µg/mL streptomycin, and 0.2 mM luciferin; pH 7.2] containing various concentrations of compounds (final 0.2% DMSO), and luminescence was recorded every 30 mins for 5 days in a luminometer LumiCEC (Churitsu).

### Protein Thermal Shift Assay

CRY1(PHR) and CRY2(PHR) recombinant proteins^40^ were diluted to 2 µM with DSF buffer (20 mM HEPES-NaOH, 150 mM NaCl, 2 mM DTT; pH 7.5) and dispensed into a 384-well white PCR plate (Bio-Rad, Cat. #MSP3852) at 17 µL per well, followed by the application of 1 µL of compounds (final 5% DMSO). The mixtures were incubated at room temperature with gentle shaking for 60 mins. 2 µL of 50X SYPRO Orange in DSF buffer (final 5X SYPRO Orange) was added, and thermal denaturation was performed using a real-time PCR detection system, CFX384 Touch (Bio-Rad).

### Quantification

For reporter activity repression, the half maximal inhibitory concentrations (IC_50_ or log[IC_50_]) were obtained by sigmoidal dose-response fitting of dilution series data with Prism software (GraphPad Software). Circadian period was determined from luminescence rhythms by a curve fitting program MultiCycle (Actimetrics). Data from the first day was excluded from analysis, because of transient changes in luminescence upon medium exchange. Concentrations for 4 hrs period-lengthening (EC_4h_) were obtained by exponential growth fitting of dilution series data with Prism software. In thermal shift assays, the first derivative of the fluorescence intensity was plotted as a function of temperature (dF/d*T*), and the highest peak of the curve was defined as the melting temperature. The half maximal effective concentrations (EC_50_ or log[EC_50_]) were obtained by sigmoidal dose-response fitting of dilution series data with Prism software.

### Intracranial Tumor Formation

All mouse procedures were performed under animal protocols that were approved by the Institutional Animal Care and Use Committees at University of Southern California, Children’s Hospital Los Angeles, and the University of Pittsburgh. Intracranial implantation of GSCs into 8-week-old female NOD.Cg-Prkdc^scid^ ll2rg^tm1Wjl^/SzJ (NSG, The Jackson Laboratory) mice were performed via stereotactic method and 20,000 cells were injected 2 mm posterior to bregma, 1 mm laterally, and 2 mm deep.

### *In Vivo* SHP1705 Administration

7 days following cell implantation, mice were randomized and divided into treatment groups of 1% CMC vehicle control, 10 mg/kg SHP1705, or 30 mg/kg SHP1705. The reagents were administered via oral gavage once a day, every morning until the manifestation of neurological signs (i.e., hunched back, severe weight loss, ataxia, and spinning). BLI Imaging. Mice were injected intraperitoneally with 200 μL of Beetle Luciferin Potassium Salt (5 mg/mL dissolved in PBS, Promega, Cat. #E1605). Mice were anesthetized with isoflurane (5%, 100% oxygen). 15 mins after injection, mice were placed in the IVIS Spectrum BLI system for imaging and anesthesia was maintained with isoflurane (2.5%, 100% oxygen). Luminescence was quantified with Living Image and data was plotted in Excel. Statistical analysis was performed with two-way ANOVA.

Survival Analysis. Once mice began to experience neurological symptoms (i.e. hunched back, significant weight loss, circling, etc.) Survival data was plotted, and statistical analysis was performed with log-rank (Mantel-Cox) test in GraphPad Prism.

### SHP1705 and SR29065 Synergy Assays

Cell viability assays were conducted as mentioned above. Relative percentages of cell viability were plotted into Excel to generate a table of SHP1705 vs. SR29065 treatments. Excel sheets were uploaded onto the web application SynergyFinder3.0 (https://synergyfinder.fimm.fi/) to visualize dose-response data and calculate the synergy score.^26^ Detection of outliers and LL4 curve fitting was performed by the program.

## RESULTS

### CRY2 is elevated in GBM tissues compared to non-cancerous samples

We previously reported the anti-GSC effects of the CRY2-selective compound SHP1703, which showed improved efficacy compared to KL001, which does not display a CRY isoform selectivity.^14,15,17^ To interrogate the molecular basis for this result, we compared clock gene levels in GBM tissue compared to non-tumor tissue in the Cancer Genome Atlas (TCGA) and CCGA datasets. We found that *CRY2* mRNA levels are decreased in GBM tissue while *CRY1* is increased **(Fig. 1B, Sup. Fig. 2A**). The differences in *CRY2* and *CRY1* levels could explain the increased efficacy of SHP1703 compared to KL001 against GSCs and provide a rationale for CRY2-selective modulation in GBM. We \hypothesized that stabilizing and elevating CRY2 protein levels could improve the ability of CRY2 to inhibit BMAL1-CLOCK transcriptional activity in GSCs and have a greater anti-GSC effect compared to targeting CRY1.

### Phase 1 study indicates that SHP1705 is safe and well-tolerated in humans

To further improve upon the efficacy and bioavailability of existing CRY2-targeting compounds we developed a novel scaffold, SHP1705 **(Sup. Fig. 1A)**, based on SHP656 and SHP1703, for oral administration to patients.^17^

We conducted a Phase I randomized, double-blind, placebo-controlled study with healthy human volunteers. In a single ascending dose (SAD) phase of the trial, a total of 56 subjects were randomized and treated, with 42 receiving single oral doses of SHP1705 and 14 receiving the placebo. In the multiple ascending dose (MAD) phase, a total of 16 subjects were randomized and treated, with 12 receiving multiple oral doses of SHP1705 and 4 receiving placebos. Both single and multiple doses of SHP1705 were well tolerated in this study. In the SAD portion of the study, the most frequently reported Treatment-Emergent Adverse Events (TEAE) overall was headache, which occurred in 5 subjects who received SHP1705 and 2 subjects who received placebo. Diarrhea and nausea were reported by 3 subjects who received SHP1705. In the MAD portion of the study, 3 subjects who received SHP1705 had reported TEAEs and none of the TEAEs were reported by more than one subject each. Subject 2205 in the Treatment I group (25 mg SHP1705 tablets, fed) discontinued the study early because of a TEAE of ventricular extrasystoles, which was considered unrelated to the study drug. There were no apparent trends in the TEAE frequency or type by dose, dose frequency (single or multiple doses), formulation (tablet or capsule), fed or fasted state, or time of dose (morning or evening). Likewise, there were no treatment related trends noted for changes in clinical laboratory parameters, vital signs, physical examinations, or electrocardiograms. The summary of single dose SHP1705 pharmacokinetic (PK) parameters were measured for 500 mg oral treatment for the max concentration of SHP1705 (C_max_), time of maximal SHP1705 (T_max_), exposure to the SHP1705 (AUC_0-t_), and half-life of SHP1705 (T_1/2_) **(Sup. Fig. 1B)**.

Analysis of biomarker data showed that expression of specific BMAL1-CLOCK target genes, *PER1* and D site albumin promoter Binding Protein (*DBP*), were altered at the highest dosage of SHP1705 (500 mg) administration in the morning, supporting target engagement **(Sup. Fig. 1C)**. Expression levels of non-circadian, housekeeping genes were not altered. **(Sup. Fig. 1D)** The blood transcriptome is partially under circadian control and can therefore provide information about circadian rhythmicity in the SCN and peripheral clocks.^19–22^ Blood transcriptome-based biomarkers genes **(Sup. Fig. 1E)** were generally not significantly altered by SHP1705 dosing, although some trends may have been masked by high variability in the data.

### SHP1705 displays significantly improved anti-GSC effects compared to earlier CRY activator scaffolds

We treated numerous GSCs with differences in TMZ sensitivity **(Sup. Table 1)** and found that SHP1705 is more efficacious against GSCs compared to non-malignant (NM) neuronal cultures, indicating potential for a therapeutic window. When the GSCs were differentiated (DGCs), they lost their sensitivity to SHP1705 treatment which further supports previous reports that it is truly the cancer stem cell population that is susceptible to clock perturbation **(Fig. 1C)**.

Loss of stemness was indicated by lack of neurosphere formation and plate-adherence **(Sup. Fig. 2B)**. We treated multiple GSCs, DGCs, and NM cells for two different days before accessing cell viability given that cell lines either respond to circadian clock compound treatment at 3 or 4 days. SHP1705 generally displayed increased efficacy against GSCs compared to earlier scaffolds KL001, SHP656, and SHP1703, with the TMZ-sensitive line MGG 31 showing the highest increased sensitivity to SHP1705 out of all the GSCs tested **(Fig. 1C-G, Sup. Fig. 2C-G)**.

### SHP1705 modulates circadian gene expression and output

To confirm increased repression of BMAL1-CLOCK transcriptional activity, we examined expression of CCGs following SHP1705 treatment. SHP1705 significantly decreased expression of *PER2*, which is under direct control of BMAL1-CLOCK **(Fig. 2A)** in GSCs compared to non-cancerous NM cells following 24hrs of treatment. Treatment of SHP1705 decreased cell counts and neurosphere size and count in GSCs compared to NM290 **(Sup. Fig. 3)**. We assessed circadian gene cycling in osteosarcoma U2OS luciferase reporter cells. Dosages were chosen starting at the concentration that reduced GSC viability to 50% (IC_50_) **(Fig. 1C, 1G)**. We found that SHP1705 displayed period lengthening and arrhythmicity in higher concentrations in the U2OS *Bmal1-dLuc* cells **(Fig. 2B)**. SHP1705 reduced amplitude of the luciferase reporter at even the lowest doses of treatment and caused arrhythmicity in the remaining dosages in *Per2-dLuc* reporter cells **(Fig. 2C)**. These results indicate that SHP1705 represses GSC growth and BMAL1-CLOCK activity and, consequently, CCG output.

**Figure 2.**
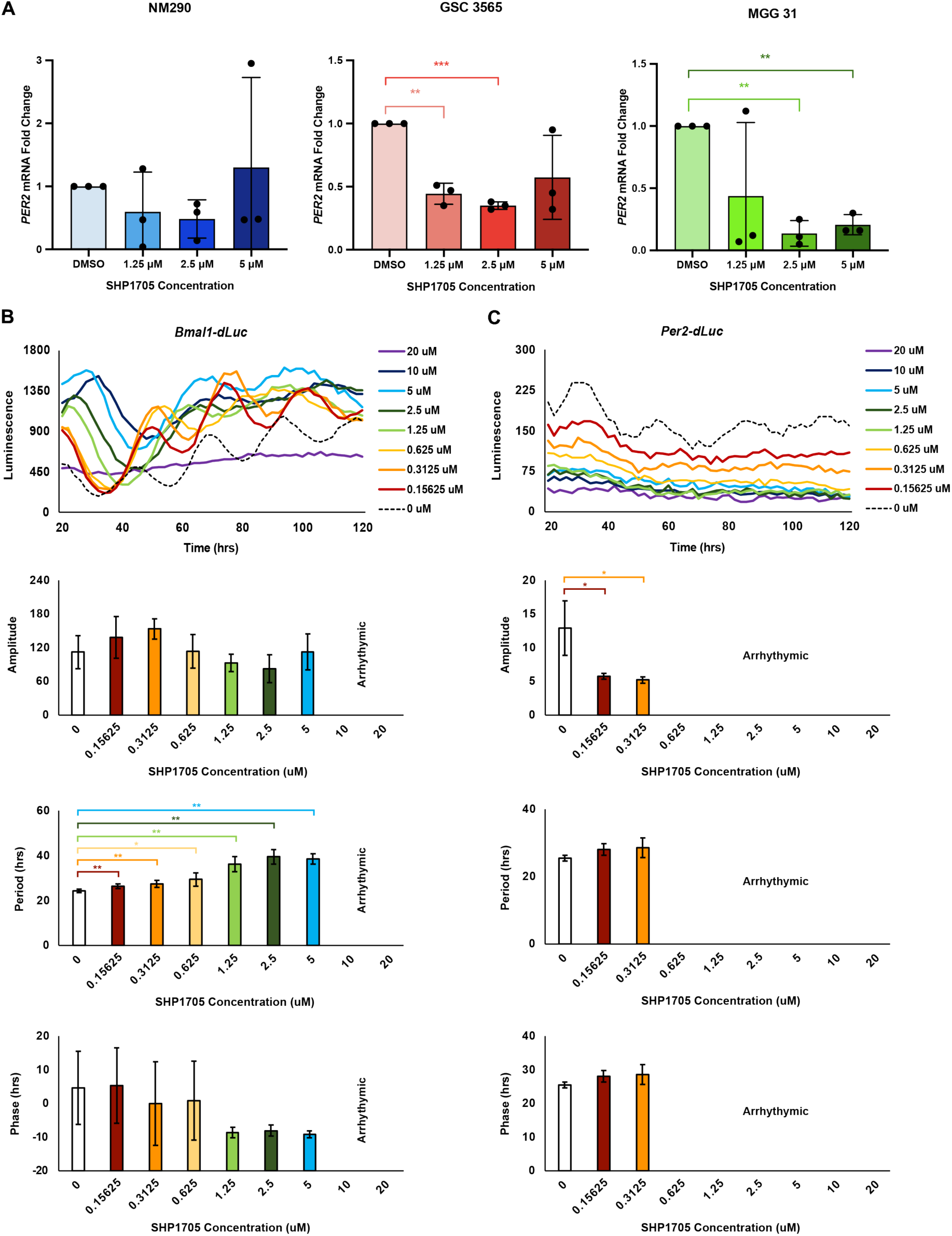
SHP1705 directly modulates circadian gene expression. **(A)** *PER2* mRNA expression in NM 290, GSC 3565, and MGG 31 was measured by qPCR following 24 hrs of DMSO control or SHP1705 treatment at 1.25, 2.5, or 5 μM (*n* = 3 biologically independent samples). Fold change was normalized to the *18S rRNA* house-keeping gene and then DMSO control. Statistical analysis was performed with two-way ANOVA. * p < 0.05, ** p < 0.01, *** p < 0.001 **(B)** U2OS *Bmal1-dLuc* or **(C)** *Per2-dLuc* reporter cells were treated with increasing doses of SHP1705 (*n* = 3-4 biologically independent samples). Luminescence was read every 2 hrs for 120 hrs. Luciferase intensity (amplitude), period, and phase was measured and plotted below Statistical analysis was performed with two-way ANOVA * p < 0.05, ** p < 0.01, *** p < 0.001

### SHP1705 is selective for the CRY2 isoform

We next aimed to confirm SHP1705’s selectivity for CRY2, like its predecessors SHP656 and SHP1703.^15^ We analyzed the effects of KL001, SHP656, SHP1703, and SHP1705 on endogenous CRY1 and CRY2 activity to repress *Per2* expression in fibroblasts from *Per2::Luc* knock-in mice. SHP1705 repressed *Per2::Luc* reporter activity in wild type cells containing both *Cry1/2* to a greater extent than either KL001, SHP656, or SHP1703 **(Fig. 3A *Top Left*, 3B)**. There was no repression of reporter activity in *Cry1/2* double knockout cells suggesting that the effects of all the tested CRY activators are indeed CRY-dependent **(Fig. 3A *Top Right*, 3B).** In *Cry1* knockout cells, while KL001 was less effective than intact cells, SHP1705 was still able to repress *Per2* reporter activity and to a greater extent than SHP656 or SHP1703 **(Fig. 3A *Bottom Left*, 3B)**. In *Cry2* knockout cells, SHP1705, SHP656, and SHP1703 showed a reduced overall repression of *Per2*::*Luc* reporter activity compared to wild type calls, with no differences in activity between these compounds **(Fig. 3A *Bottom Right*, 3B)**. In *Cry1/2* wild type cells, SHP1705 showed a greater period lengthening effect on compared to either SHP1703 or KL001 **(Fig. 3C *Top Left*, 3D)**. SHP1705 was able to lengthen the period in *Cry1* knockout and *Cry2* knockout cells, but this was to a greater or lesser extent, respectively, compared to *Cry1/2* wild type cells **(Fig. 3C *Bottom*, 3D)**. The effects observed in *Cry1* knockout cells could be contributed to *Cry2* compensation and upregulation. We conducted a thermal shift assay to evaluate direct interaction with the photolyase homology region (PHR) of CRY1 and CRY2. We found that SHP1705 exhibited a preference for CRY2 compared to KL001 and, even, SHP1703 **(Fig. 3E-F)**. These results confirm SHP1705 activity is mediated by selectively interacting with CRY2 and highlights its increased affinity for CRY2 compared to earlier scaffolds.

**Figure 3.**
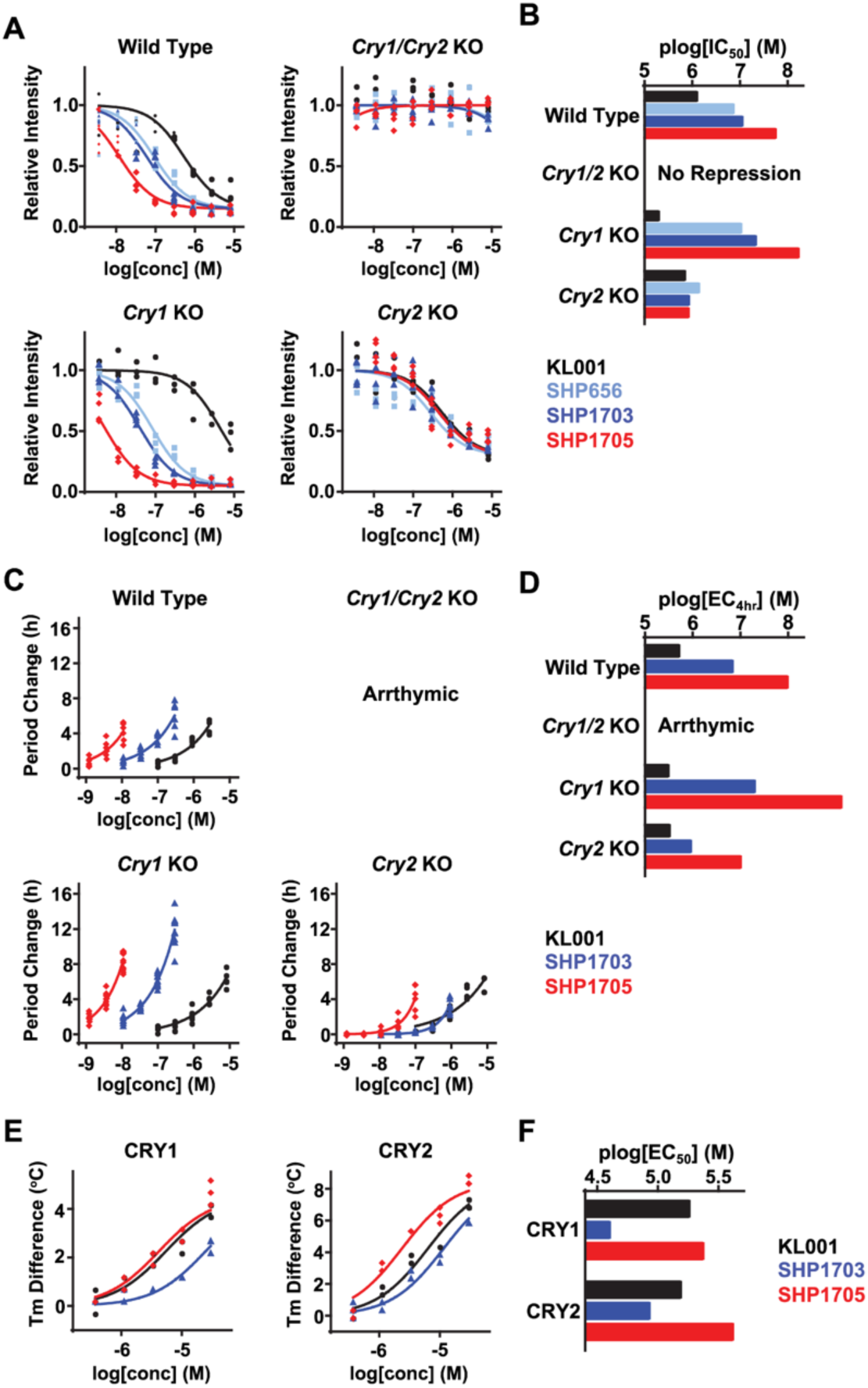
SHP1705 is selective for the CRY2. **(A)** Effects on *Per2::Luc* knock-in reporter activity in wild type, *Cry1/Cry2* double knockout, *Cry1* knockout, and *Cry2* knockout MEFs. Changes in luminescence intensity compared to DMSO control are shown (*n* = 3-4 biologically independent samples). **(B)** Concentrations for 50% inhibition (plog[IC_50_] that represents -log[IC_50_]) are plotted. **(C)** Effects on circadian period in *Per2::Luc* knock-in MEFs. Changes in period compared to DMSO control are shown (*n* = 4-12 biologically independent samples). **(D)** Concentrations of a 4 hr period lengthening (plog[EC_4hr_] that represents -log[EC_4hr_]) are plotted. **(E)** Interaction with CRY1(PHR) and CRY2(PHR) *in vitro*. Changes in denaturing temperatures of recombinant CRY(PHR) proteins in the presence of various concentrations of compounds compared to DMSO control are shown (*n* = 3 biologically independent samples). **(F)** Concentrations for 50% stabilization (plog[EC_50_] that represents -log[EC_50_]) are plotted.

### SHP1705 delays GBM tumor growth rate and increases survival

To interrogate the effects of SHP1705 against GBM tumors, we established GBM patient-derived xenografts (PDX) initiated using different GSCs **(Fig. 4A)**. SHP1705 was well-tolerated and did not appear to disrupt overall circadian function as there was no evidence of significant weight gain or loss **(Sup. Fig. 4)**. Use of hGBM18 Firefly luciferase mCherry (FMC) Mouse Passaged 1 (MP1) cells allowed for visual monitoring of tumor growth via bioluminescence imaging (BLI) and showed that, although not statistically significant, SHP1705 had a trend of decreasing the tumor growth rate in a dose-dependent manner compared to vehicle control **(Fig. 4B)**. In mice bearing tumors formed from either T387 GSC or T3565 GSC patient-derived lines transduced with a luciferase reporter, treatment with SHP1705 conferred longer survival times compared to mice given vehicle control **(Fig. 4C-D)**. Mice treated with SHP1705 survived longer than those treated with SHP656 **(Fig. 4D)**. These results provide further support for CRY2 activators as a novel paradigm for GBM treatment and SHP1705’s superior *in vivo* efficacy compared to earlier scaffolds.

**Figure 4.**
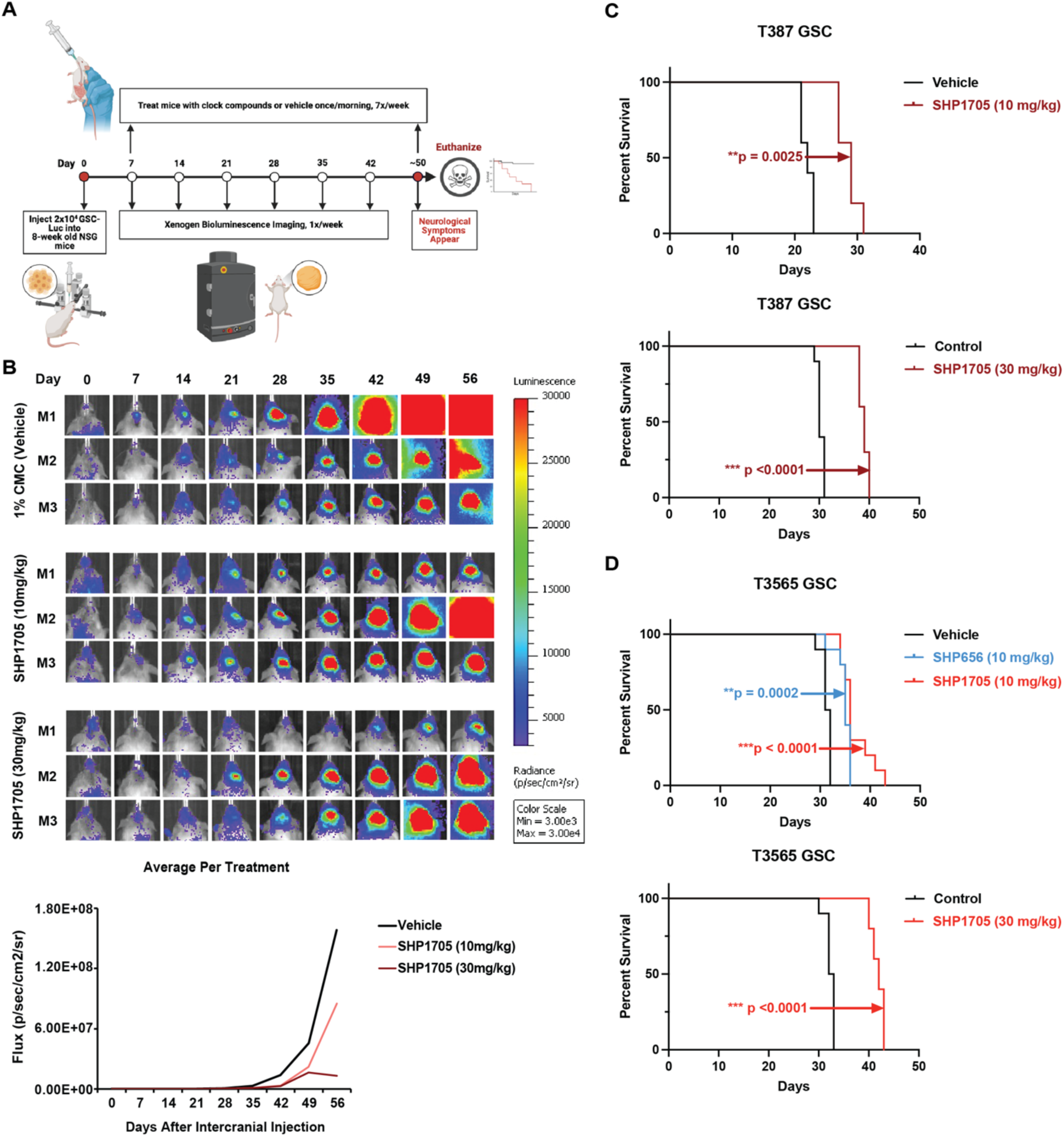
SHP1705 delays GBM tumor growth rate and increases survival. **(A)** *in vivo* pipeline for GBM PDX tumor establishment. 2×10^4^ GSCs containing a constitutive luciferase reporter are injected into 8-week-old NSG mice. 7 days after the surgery, mice are imaged with the Xenogen BLI system and are imaged thereafter once a week. Mice are randomized into treatment groups based on the 7 day-post-surgery luciferase signal and given vehicle control or SHP1705 once every morning, every day. Mice are euthanized once neurological symptoms appear. Image was created with BioRender. **(B)** Top: Bioluminescence images of representative mice injected with hGBM18 FMC MP1 and treated with for 1% CMC vehicle control, SHP1705 (10 mg/kg), and SHP1705 (30 mg/kg). Bottom: Quantification of radiance (p/sec/cm^2^/sr) (*n* = 10-11 mice/group) indicates size of tumor as a function of luciferase signal. **(C)** Kaplan-Meier survival curves of GBM PDX tumors established with T387 GSCs and treated with 1% CMC vehicle or 10 or 30 mg/kg of SHP1705. Statistical analysis was performed with log-rank (Mantel-Cox) test. **(D)** Kaplan-Meier survival curves of GBM PDX tumors established with T3565 GSCs and treated with 1% CMC vehicle, 10 mg/kg of SHP656, or 10 or 30 mg/kg of SHP1705 (*n* = 5 mice/group). Statistical analysis was performed with log-rank (Mantel-Cox) test.

### SHP1705 synergizes with novel REV-ERB agonist SR29065

Given that monotherapy treatment with SHP1705 or combination treatment using earlier REV-ERB agonists and CRY activators were effective against GSCs, we decided to interrogate the combination of SHP1705 with a novel REV-ERB agonist, SR29065.^7,23^ REV-ERB agonists promote REV-ERBs’ suppression of *BMAL1* transcription by increasing engagement of REV-ERBαβ with the Nuclear CoRepressor (NCoR) and Histone Deacetylase 3 (HDAC3) **(Fig. 5A)**.^12^ SR29065 is a novel structure that is not related to earlier REV-ERB agonists SR9009 and SR9011, which have been shown to have REV-ERB independent effects.^24^ SR29065 is selective for REV-ERBα and has demonstrated to provide therapeutic benefit in a mouse model of autoimmune disorders.^23^ We first tested SR29065 as a monotherapy against GSCs. Not only did SR29065 display specific effects against GSCs compared to NM cells and DGCs, like CRY activators, but it also had lower IC_50_’s against GSCs compared to SR9009 and SR9011 **(Fig. 5B-E, Sup. Fig. 5A-D**). SR29065 altered the amplitude and resulted in arrhythmicity at higher doses in our luciferase reporter activity in U2OS *Bmal1-dLuc* cells, indicating it indeed affects CCG output **(Fig. 5F)**. Although not statistically significant, SR29065 decreased the amplitude of luciferase intensity at higher dosages of treatment in the U2OS *Per2-*dLuc reporter cells **(Fig. 5G)**. Overall, CRY activators were found to reduce *Per2* **(Fig. 2C)** intensity while REV-ERB agonists reduces *Bmal1* intensity of the luciferase reporters **(Fig. 5F)**, as predicted from their mechanism of action.

**Figure 5.**
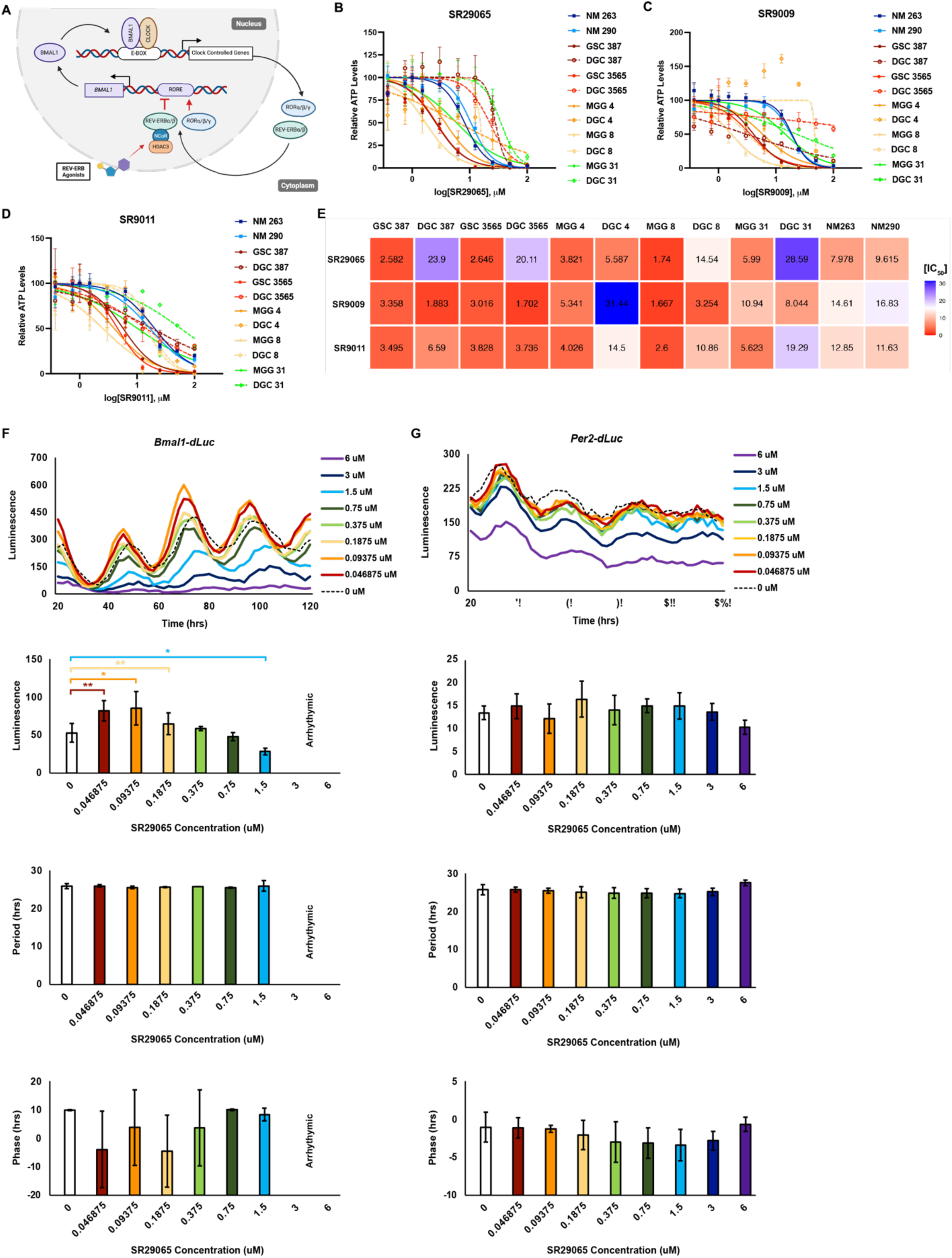
Novel REV-ERB agonist SR29065 has increased potency against GSCs. **(A)** Mechanism of REV-ERB agonists. REV-ERB agonists increase binding of REV-ERBαβ with their binding partners, NCoR and HDAC3, to increase repression of *BMAL1* transcription. **(B-E)** Cell viability following **(B)** SR29065, **(C)** SR9009, or **(D)** SR9011 treatment for 3 days. **(E)** Heatmap of summarized IC_50_ values following CRY activator treatment in control NM cells, GSCs, and DGCs. **(F)** U2OS *Bmal1-dLuc* or **(G)** *Per2-dLuc* reporter cells were treated with increasing doses of SR29065 (*n* = 4 biologically independent samples). Luminescence was read every 2 hrs for 120 hrs. Luciferase intensity (amplitude), period, and phase was measured and plotted below. Statistical analysis was performed with two-way ANOVA * p < 0.05, ** p < 0.01, *** p < 0.001

We then combined SHP1705 and SR29065 and treated several matched GSC and DGC lines with the combination to generate synergy plots and zero interaction potency (ZIP) synergy scores for each cell line. The ZIP scoring model is used to determine the change in potency of dose-response curves between single drugs and drug combinations. A synergy score that is less than -10 means that the two drugs are antagonistic. One that is from -10 to 10, the two drugs are likely to be additive but the closer the score is to 0, the less confidence there is with regards to synergy or antagonism. Lastly a score that greater than 10 means that the two drugs are synergistic.^25,26^ Most of the matched GSC and DGC cell lines showed synergy when the two compounds were combined. **(Fig. 6A-E)**. However, the combined treatment did not display synergistic effects against DGC 4, or against either the TMZ resistant line MGG 31 or its differentiated counterpart DGC 31 **(Fig. 6F-H)**. This suggests that either monotherapy with either SHP1705 or SR29065 alone in these cells is sufficient to cause anti-GSC effects or the high sensitivity to SR29065 combined with SHP1705 yields a synergistic effect that is occurring earlier than 3 days and the remaining analyzed cells are resistant populations. Further experiments will have to be done across a time course to capture the full synergy panel across all cell lines. These results are encouraging in that they generate a hypothesis that SHP1705 could potentiate the standard-of-care (radiation plus TMZ) in TMZ-resistant cells/tumors too. Taken together, these results demonstrate that targeting the two negative limbs of the circadian clock can provide combination effects against GSCs and differentiated GBM cells.

**Figure 6.**
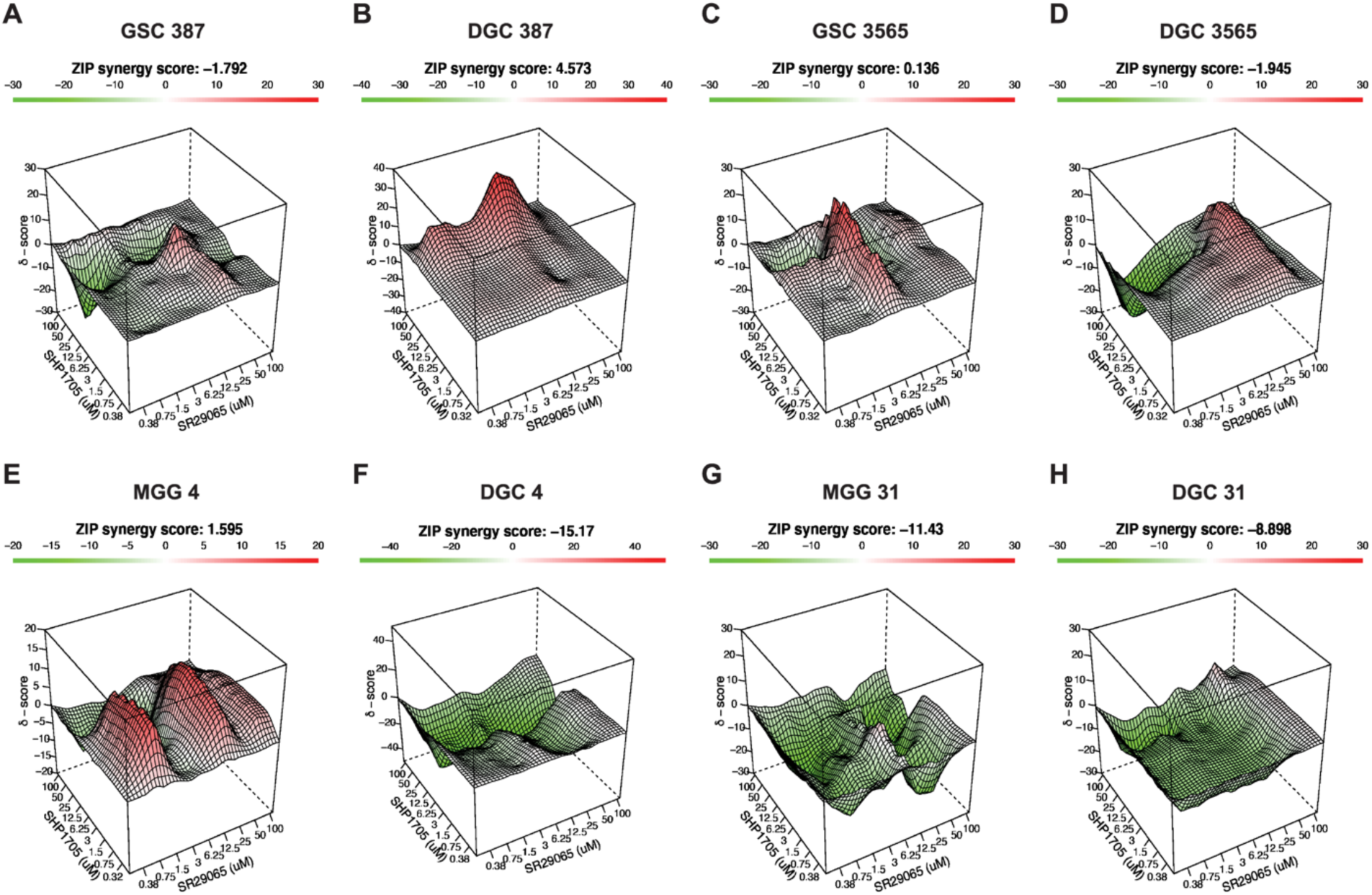
SHP1705 synergizes with novel REV-ERB agonist SR29065. **(A)** GSC 387, **(B)** DGC 387, **(C)** GSC 3565, **(D)** DGC 3565, **(E)** MGG 4, **(F)** DGC 4, **(G)** MGG 31, or **(H)** DGC 31 was treated with both SHP1705 and SR29065 for 4 days (*n* = 3-4 biologically independent samples). Cell viability results were plotted with SynergyFinder3.0.

## DISCUSSION

The current state of GBM treatment is dismal, with treatments after tumor recurrence only offering mostly palliative care. Patients who receive only surgical resection have a median survival of 3 to 6 months. The addition of radiation increases median survival to 12 months and, the current standard of care of radiation with concomitant TMZ (chemoradiation) still only offers a median survival of 14-16 months.^27^ GBM is highly infiltrative meaning surgeons must balance both maximal surgical resection without compromising neurological functions.^28^ Additionally, the presence of GBM stem cells makes treatment even more challenging due to their existence well beyond tumor margins and resistance to chemoradiation amongst their ability to initiate new tumors, secrete angiogenic factors, and promote an immune-suppressive environment by increasing microglia infiltration.^5,11^

The circadian clock has emerged in recent years as a novel therapeutic target in GBM that specifically addresses the survival of the GSC population as well targeting the tumor-supporting environment.^7–9,11,13^ The core clock protein BMAL1 has an increased chromatin-binding pattern that overlaps with marks of active transcription in GSCs compared to noncancerous neural stem cells. This increased clock-mediated transcriptional control, which is not observed in other GBM or neural-related cells, may allow targeting for targeting of clock components.^7^

We probed this hypothesis by investigating the novel CRY activator SHP1705, which has been found to be safe and well-tolerated in healthy humans in Phase 1 trials **(Sup. Fig. 1B-D)**. In comparison to non-malignant (NM) neuronal cells, the GSCs were much more susceptible to SHP1705 treatment. This sensitivity is independent of TMZ responsiveness. Remarkably, once the GSCs were allowed to differentiate (DGCs), they lost their sensitivity to SHP1705 **(Fig. 1C, Sup. Fig. 2C)**. These results suggest that, like genetic perturbation, pharmacological targeting of BMAL1-CLOCK transcriptional output selectivity attenuates GSC survival and is not observed in non-stem cell cancer cells nor non-cancerous cells. This further supports our hypothesis and other reports that have demonstrated that GSCs are uniquely reliant on the clock-controlled transcription. Furthermore, this novel CRY activator scaffold has increased potency compared to earlier scaffolds, KL001, SHP656, and SHP1703 **(Fig. 1C-G, Sup. Fig. 2C-F)**.

To confirm the mechanism of action of SHP1705, we examined luciferase reporter activity following treatment of SHP1705. SHP1705 treatment resulted in increased repression of BMAL1-CLOCK transcriptional activity. Treatment of SHP1705 decreased the expression of BMAL1-CLOCK E-box target gene, *PER2*, in GSCs and had minimal effects on noncancerous NM cells **(Fig. 2A)**. This effect was observed directly through the dose-dependent decrease in intensity of the luciferase levels in the *Per2-dLuc* cells **(Fig. 2B)**. It was observed indirectly through the increase in the intensity of the luciferase reporter and elongation of periodicity of the luciferase reporter rhythms in the *Bmal1-dLuc* cells, which suggests that *Nr1d1/2* (REV-ERBs) expression is reduced, resulting in decreased repression of *Bmal1* transcription **(Fig. 2C**).

We characterized the preferential interaction of SHP1705 with CRY2. Through a series of cell-based experiments including *Per2::Luc* luciferase assays and protein thermal shift assays, we found that like its predecessors, SHP1705 has isoform selectivity for CRY2 over CRY1. Furthermore, SHP1705 displays even stronger preference for CRY2 than SHP656 and SHP1703 **(Fig. 3)**. Because *CRY2* mRNA levels are decreased in GBM tumors compared to normal tissue, specifically stabilizing CRY2 protein levels over CRY1 may prove to be especially beneficial for GBM treatment **(Fig. 1B, Sup. Fig. 1A)**.

We show in this study that SHP1705 treatment *in vivo* reduces the tumor growth rate of GBM PDX tumors and increases survival time of our mouse model **(Fig. 4)**. The lack of significant weight gain or loss in low and higher doses of SHP1705 along with the Phase 1 studies further supports that SHP1705 does not result in cytotoxicity. SHP1705 did not adversely disrupt overall circadian rhythms under the light-dark cycle condition as indicated by the lack of significant weight gain **(Sup. Fig. 1B-D & 4)**.^29,30^

REV-ERB agonists have previously been reported to show anti-GSC efficacy. SHP1705’s single agent efficacy predicts that combining CRY activators with a REV-ERB agonist might yield synergistic effects on the circadian clock TTFL by suppressing further *BMAL1* transcription and, consequently, GSC viability. We probed the novel REV-ERB agonist SR29065 in combination with SHP1705 to determine whether targeting both feedback loops of the clock will result in synergistic effects against GSCs.^7^ SR29065 displayed increase efficacy against GSCs compared to earlier REV-ERB agonist scaffolds, SR9009 and SR9011. Like SHP1705, SR29065 displays selectivity against GSCs over normal neuronal cells and DGCs **(Fig, 5B-D, Sup. Fig. 5)**. In luciferase reporter assays, SR29065 displayed its mechanism-of-action in increasing REV-ERB-mediated repression of *Bmal1* transcription through the observed decrease in luciferase intensity in both *Bmal1*-dLuc and *Per2*-dLuc reporter cells and the periodicity elongation in *Bmal1-dLuc* cells **(Fig. 5E-F)**. It was previously shown that SR29065 reduced REV-ERBα target genes *Bmal1* and *Clock* in mice livers following treatment compared to vehicle.^23^ We found that combinations of SHP1705 and SR29065 displayed synergistic effects in multiple GSC and DGC lines, supporting our hypothesis that simultaneously targeting both feedback loops of the clock potentiate clock-mediated anti-GBM effects **(Fig. 6A-E)**. However, some GSC and DGC cell lines did not show synergistic effects, most notably in a TMZ-resistant cell line **(Fig. 6F-H)**. Combined dosing may support the growth of clock compound treatment-resistant subpopulations. Combining either clock compound with TMZ and radiation may sensitize these cells to standard-of-care therapy. Since we see that differentiated GSCs lose their specific sensitivity to clock modulation, it will likely be important to combine any potential clock-targeting therapy with radiation and TMZ, which will select for the sensitive GSC population.

These results highlight the single agent efficacy of SHP1705 and that can be combined with other clock compounds, such as SR29065, or even chemoradiation as a novel GBM treatment paradigm that targets GSCs. As GSCs present a major obstacle in the treatment of GBM by promoting tumor recurrence, clock perturbation via the administration of orally bioavailable, well-tolerated compounds can provide an easily accessible yet effective treatment option for patients with minimal side effects. With increased understanding of how response to chemoradiation changes according to time-of-day administration, we can take advantage of different modalities of chronomedicine to improve upon GBM patient outcomes.^13,31–34^ We can do so by combining two methods: targeting circadian clock components directly and timing dosing of treatments to align with the maximal effects. BMAL1-CLOCK are required not only for cell autonomous survival and stemness of GSCs but also play an integral part in angiogenesis and immunosuppression in the GBM TME.^7–9^ Therefore, targeting the clock has multiple beneficial effects in GBM treatment and this now needs to be extensively explored in combination treatment paradigms with current standard of care. Beyond this, chronomedicine can be utilized to widen personalized and precision-based medicine that is currently heavily reliant on genetic testing, which is not yet widely available nor affordable for all patient populations.

## AUTHORSHIP

P.C., S.A.K, and J.N.R. conceived the project and main conceptual ideas. P.C., Y.N., and T.H. designed and carried out the *in vitro* experiments. P.C., Q.W., A.H. S.M., G.K., and R.A.M. designed and carried out the animal experiments. J.C. provided SHP1705 and supervised the Phase 1 study. J.W. generated R script for heatmaps. T.K. and L.A.S. synthesized SR29065 and provided experimental methods for the compound. S.A.K. and J.N.R. supervised the findings of the work. P.C. and S.A.K. wrote the manuscript. All authors discussed the results and contributed to the final version of the manuscript.

## DATA AVAILABILITY

All data will be made available upon request to the corresponding author.

## ACKNOWLEDGMENTS

We would like to thank Khalid Shah and Hiroaki Wakimoto for sharing the MGG and hGBM18 FMC MP1 cell lines.

## REFERENCES

1. Schaff, L. R. & Mellinghoff, I. K. Glioblastoma and Other Primary Brain Malignancies in Adults: A Review. JAMA 329, 574–587 (2023).

2. Mohammed, S., Dinesan, M., Ajayakumar, T. Survival and quality of life analysis in glioblastoma multiforme with adjuvant chemoradiotherapy: a retrospective study. Rep. Pract. Oncol. Radiother. 27, 1026–1036 (2022).

3. Stupp, R. Hegi, M. E., Mason, W. P., van den Bent, M. J., Taphoorn, M. J. B., Janzer, R. C., et al. Effects of radiotherapy with concomitant and adjuvant temozolomide versus radiotherapy alone on survival in glioblastoma in a randomised phase III study: 5-year analysis of the EORTC-NCIC trial. Lancet Oncol. 10, 459–466 (2009).

4. Lathia, J. D., Mack, S. C., Mulkearns-Hubert, E. E., Valentim, C. L. L., Rich, J. N. Cancer stem cells in glioblastoma. Genes Dev. 29, 1203–1217 (2015).

5. Gimple, R. C., Bhargava, S., Dixit, D., Rich, J. N. Glioblastoma stem cells: lessons from the tumor hierarchy in a lethal cancer. Genes Dev. 33, 591–609 (2019).

6. Prager, B. C., Bhargava, S., Mahadev, V., Hubert, C. G., Rich, J. N. Glioblastoma Stem Cells: Driving Resiliency through Chaos. Trends Cancer 6, 223–235 (2020).

7. Dong, Z., Zhang, G., Qu, M., Gimple, R. C., Wu, Q., Qiu, Z. et al. Targeting Glioblastoma Stem Cells through Disruption of the Circadian Clock. Cancer Discov. (2019) doi:10.1158/2159-8290.CD-19-0215.

8. Chen, P., Hsu, W-H., Chang, A., Tan, Z., Lan, Z., Zhou, A. et al. Circadian Regulator CLOCK Recruits Immune-Suppressive Microglia into the GBM Tumor Microenvironment. Cancer Discov. 10, 371–381 (2020).

9. Pang, L., Dunterman, M., Xuan, W., Gonzalez, A., Lin, Y., Hsu, W-H. et al. Circadian regulator CLOCK promotes tumor angiogenesis in glioblastoma. Cell Rep. 42, 112127 (2023).

10. Takahashi, J. S. Transcriptional architecture of the mammalian circadian clock. Nat. Rev. Genet. 18, 164–179 (2017).

11. Chan, P., Rich, J. N., Kay, S. A. Watching the clock in glioblastoma. Neuro-Oncol. 25, 1932– 1946 (2023).

12. Solt, L. A. Wang, Y., Banerjee, S., Hughes, T., Kojetin, D. J., Lundasen, T., et al. Regulation of circadian behaviour and metabolism by synthetic REV-ERB agonists. Nature 485, 62–68 (2012).

13. Battaglin, F., Chan, P., Pan, Y., Soni, S., Qu, M., Spiller, E. R. et al. Clocking cancer: the circadian clock as a target in cancer therapy. Oncogene 1–14 (2021) doi:10.1038/s41388-021-01778-6.

14. Hirota, T., Lee, J. W., St. John, P. C., Sawa, M., Iwaisako, K., Noguchi, T., et al. Identification of small molecule activators of cryptochrome. Science 337, 1094–1097 (2012).

15. Miller, S., Kesherwani, M., Chan, P., Nagai, Y., Yagi, M., Cope, J. et al. CRY2 isoform selectivity of a circadian clock modulator with antiglioblastoma efficacy. Proc. Natl. Acad. Sci. 119, e2203936119 (2022).

16. Humphries, P. S., Bersot, R., Kincaid, J., Mabery, E., McCluskie, K., Park, T. et al. Carbazole-containing sulfonamides and sulfamides: Discovery of cryptochrome modulators as antidiabetic agents. Bioorg. Med. Chem. Lett. 26, 757–760 (2016).

17. Humphries, P. S., Bersot, R., Kincaid, J., Mabery, E., McCluskie, K., Park, T. et al. Carbazole-containing amides and ureas: Discovery of cryptochrome modulators as antihyperglycemic agents. Bioorg. Med. Chem. Lett. 28, 293–297 (2018).

18. Fan, W., Caiyan, L., Ling, Z., Jiayun, Z. Aberrant rhythmic expression of cryptochrome2 regulates the radiosensitivity of rat gliomas. Oncotarget 8, 77809–77818 (2017).

19. Hughey, J. J. Machine learning identifies a compact gene set for monitoring the circadian clock in human blood. Genome Med. 9, 19 (2017).

20. Laing, E. E., Möller-Levet, C. S., Poh, N., Santhi, N., Archer, S.N., Dijk, D-J. Blood transcriptome based biomarkers for human circadian phase. eLife 6, e20214 (2017).

21. Wittenbrink, N., Ananthasubramaniam, B., Münch, M., Koller, B., Maier, B., Weschke, C. et al. High-accuracy determination of internal circadian time from a single blood sample. J. Clin. Invest. 128, 3826–3839 (2018).

22. Huang, Y. & Braun, R. Platform-independent estimation of human physiological time from single blood samples. Proc. Natl. Acad. Sci. 121, e2308114120 (2024).

23. He, Y., Zhu., Greenman, K., Ruiz, C., Shang, J., Lu, Q., et al. Structure–Activity Relationship and Biological Investigation of a REV-ERBα-Selective Agonist SR-29065 (34) for Autoimmune Disorders. J. Med. Chem. 66, 14815–14823 (2023).

24. Dierickx, P., Emmett, M. J., Jiang, C., Uehara, K., Liu, M., Adlanmerini, M. et al. SR9009 has REV-ERB–independent effects on cell proliferation and metabolism. Proc. Natl. Acad. Sci. 116, 12147–12152 (2019).

25. Yadav, B., Wennerberg, K., Aittokallio, T., Tang, J. Searching for Drug Synergy in Complex Dose–Response Landscapes Using an Interaction Potency Model. Comput. Struct. Biotechnol. J. 13, 504–513 (2015).

26. Ianevski, A., Giri, A. K. & Aittokallio, T. SynergyFinder 3.0: an interactive analysis and consensus interpretation of multi-drug synergies across multiple samples. Nucleic Acids Res. 50, W739–W743 (2022).

27. Marko, N. F., Weil, R. J., Schroeder, J., Lang, F. F., Suki, D., Sawaya, R. E. Extent of Resection of Glioblastoma Revisited: Personalized Survival Modeling Facilitates More Accurate Survival Prediction and Supports a Maximum-Safe-Resection Approach to Surgery. J. Clin. Oncol. 32, 774–782 (2014).

28. Seker-Polat, F., Pinarbasi Degirmenci, N., Solaroglu, I., Bagci-Onder, T. Tumor Cell Infiltration into the Brain in Glioblastoma: From Mechanisms to Clinical Perspectives. Cancers 14, 443 (2022).

29. Salgado-Delgado, R. C., Saderi, N., del Carmen Basualdo, M., Guerro-Vargas, N. N., Escobar, C., Buijs, R. M. Shift Work or Food Intake during the Rest Phase Promotes Metabolic Disruption and Desynchrony of Liver Genes in Male Rats. PLOS ONE 8, e60052 (2013).

30. Crosby, P., Hamnett, R., Putker, M., Hoyle, N. P., Reed, M., Karam, C. J. et al. Insulin/IGF-1 Drives PERIOD Synthesis to Entrain Circadian Rhythms with Feeding Time. Cell 177, 896–909.e20 (2019).

31. Sulli, G., Manoogian, E. N. C., Taub, P. R., Panda, S. Training the Circadian Clock, Clocking the Drugs, and Drugging the Clock to Prevent, Manage, and Treat Chronic Diseases. Trends Pharmacol. Sci. 39, 812–827 (2018).

32. Damato, A. R., Luo, J., Katumba, R. G. N., Talcott, G. R., Rubin, J. B., Herzog, E. D. et al. Temozolomide chronotherapy in patients with glioblastoma: a retrospective single-institute study. Neuro-Oncol. Adv. 3, 1–11 (2021).

33. Zhanfeng, N., Yanhui, L., Zhou, F., Shaocai, H., Guangxing, L., Hechun, X. Circadian genes Per1 and Per2 increase radiosensitivity of glioma in vivo. Oncotarget 6, 9951–9958 (2015).

34. Zhu, X., Maier, G., Panda, S. Learning from circadian rhythm to transform cancer prevention, prognosis, and survivorship care. Trends Cancer 10, 196–207 (2024).

35. Zhang, G., Dong, Z., Prager, B. C., Kim, L. J. K., Wu, Q., Gimple, R. C., et al. Chromatin remodeler HELLS maintains glioma stem cells through E2F3 and MYC. JCI Insight 4, (2019).

36. Jahan, N., Lee, J. M., Shah, K., Wakimoto, H. Therapeutic targeting of chemoresistant and recurrent glioblastoma stem cells with a proapoptotic variant of oncolytic herpes simplex virus. Int. J. Cancer 141, 1671–1681 (2017).

37. Hirota, T., Lewis, W. G., Liu, A. C., Lee, J. W., Schultz, P. G., Kay, S. A. A chemical biology approach reveals period shortening of the mammalian circadian clock by specific inhibition of GSK-3β. Proc. Natl. Acad. Sci. 105, 20746–20751 (2008).

38. Zhang, E. E., Liu, A. C., Hirota, T., Miraglia, L. J., Welch, G., Pongsawakul, P. Y. et al. A Genome-wide RNAi Screen for Modifiers of the Circadian Clock in Human Cells. Cell 139, 199–210 (2009).

39. Liu, A. C., Welsh, D. K., Ko, C. H., Tran, H. G., Zhang, E. E., Priest, A. A. et al. Intercellular Coupling Confers Robustness against Mutations in the SCN Circadian Clock Network. Cell 129, 605–616 (2007).

40. Miller, S., Son, Y. L., Aikawa, Y., Makino, E., Nagai, Y., Srivastava, A. et al. Isoform-selective regulation of mammalian cryptochromes. Nat. Chem. Biol. 16, 676–685 (2020).

41. Nakao, R., Okauchi, H., Hashimoto, C., Wada, N., Oishi, K. Determination of reference genes that are independent of feeding rhythms for circadian studies of mouse metabolic tissues. Mol. Genet. Metab. 121, 190–197 (2017).

