## Supplemental Data and Materials for "Advancing Clinical Response Against Glioblastoma: Evaluating SHP1705 CRY2 Activator Efficacy in Preclinical Models and Safety in Phase I Trials"

#### SUPPLEMENTAL METHODS

**Phase 1 Study Design.** This was a first-in-human, single-center, double-blind, placebo-controlled, single- and multiple-ascending dose study of the safety, tolerability, and pharmacokinetics (PK) of oral doses of SHP1705. The study consisted of two parts: a single-ascending dose (SAD) portion, conducted in a staggered parallel sequence, and a multiple-dose (MD) portion.

In the SAD portion of the study, subjects were enrolled in seven serial cohorts of eight subjects each. Within each cohort, six subjects were assigned to receive a single dose of SHP1705 (from 25 mg in the first cohort to 500 mg in the final cohort), and two subjects were assigned to receive a single dose of placebo. Dosing within each cohort was staggered so that two of the eight subjects (i.e., sentinels) were dosed on the first day (one randomized to placebo and one randomized SHP1705). The rest of the subjects in that cohort (6 of 8) were randomized 5:1, SHP1705:placebo, and were dosed no earlier than the following day once no safety issues were noted after dosing the first two subjects. Each subject received a single dose of study medication in the morning or the evening after at least an 8-hour fast on Day 1 and remained at the research unit for a 96-hour safety and PK collection period. Subjects were discharged on Day 5. Intensive electrocardiograms (ECGs) were collected predose on Day 1 and for the first 24 hours postdose for each SAD cohort to assess the potential effect of SHP1705 on cardiac safety.

In the MD part of the study, two cohorts of eight subjects each were enrolled, randomized (6 SHP 1705:2 placebo), and dosed with study medication (SHP 1705 tablets, 25 mg) for 7 consecutive days in the morning (MAD Cohort 1) or the evening (MAD Cohort 2) after at least an 8-hour fast (Days 1-7), with a 96-hour PK collection period after the doses were administered on Days 1 and 7. Safety information was collected throughout the study. Subjects were discharged on Day 11.

**Buffy Coat Preparation.** Whole blood was drawn for plasma PK sampling within 1 hour pre-dose and at 15 mins, 30 mins, 1 hr, 2 hrs, 4 hrs, 6 hrs, 8 hrs, 12 hrs, 24 hrs, 48 hrs, 72 hrs, and 96 hrs post-dose. After plasma had been taken for PK analysis, buffy coats were prepared using standard techniques and stored in DNAse-free, RNAse-free tubes containing RNAlater (Invitrogen Cat. No. AM7020). Buffy coat aliquots were stored at 2 – 8°C within 120 minutes of blood sample collection and stored long-term at -80°C until RNA extraction. RNA extraction was performed using Trizol/chloroform extraction methods. RNA quality and concentration were measured on a Bioanalyzer 2100 according to the manufacturer's instructions. Samples with RNA Integrity Numbers (RIN) ≥ 5.0 and concentrations ≥ 1000 pg/mL were required for analysis. Samples with lower quality and/or concentration were re-extracted or newly prepared from buffy coat samples.

**cDNA preparation and qPCR.** Complementary DNA (cDNA) was synthesized from RNA samples using the High-Capacity cDNA Reverse Transcription Kit (Thermo Fisher Cat. No. 4368813). Gene expression was quantitatively measured by duplex qPCR. The primer corresponding to the target gene of interest bore the FAM™ dye, and the internal reference gene, *TBP*, bore the VIC® dye. *TBP* was selected as a reference gene since it was found to have moderate expression in buffy coat samples and is reported to be a non-rhythmic circadian control gene.[41] The oligonucleotides used as primers in this study are commercially available and are shown below.

| Gene | Dye | Catalog ID | Type | Category |
|---|---|---|---|---|
| *TBP* | VIC | Hs99999910 | m1 | Non-circadian housekeeping gene *(internal reference gene)* |
| *ARNTL (BMAL1)* | FAM | HS00154147 | m1 | Core clock |
| *CRY1* | FAM | HS00172734 | m1 | Core clock |
| *DAAM2* | FAM | HS00322497 | m1 | Blood circadian biomarker |
| *DBP* | FAM | HS00609747 | m1 | Circadian clock output gene |
| *DDIT4* | FAM | HS01111686 | g1 | Blood circadian biomarker |
| *EPHX2* | FAM | HS00932316 | m1 | Blood circadian biomarker |
| *FAAH2* | FAM | HS00415899 | m1 | Blood circadian biomarker |
| *FOSL2* | FAM | HS01050117 | m1 | Blood circadian biomarker |
| *GAPDH* | FAM | HS02758991 | g1 | Non-circadian housekeeping gene |
| *IL6* | FAM | HS00985639 | m1 | Blood circadian biomarker |
| *NRI1D1 / REV-ERBα* | FAM | HS00253876 | m1 | Core clock |
| *PER1* | FAM | HS00242988 | m1 | Core clock |
| *PER2* | FAM | HS00256143 | m1 | Core clock |
| *PPIA* | FAM | Hs04194521 | s1 | Non-circadian housekeeping gene |
| *RPLP0* | FAM | Hs00420895 | gH | Non-circadian housekeeping gene |

Each cohort contained six subjects treated with SHP-1705 (i.e., six biological replicates) and two subjects treated with placebo to match (i.e., two biological replicates). Therefore, to increase statistical power for comparison of relative expression levels after morning administration of test article, a set of six placebo biological replicates was prepared by preparing RNA from buffy coat samples isolated from the placebo-treated subjects in three cohorts. Each biological sample was analyzed by qPCR in quadruplicate (i.e. four technical replicates), arrayed on 96-well qPCR plates. For each sample, raw data (including cycle threshold (CT) values) were exported from QuantStudio 6 for analysis in Microsoft Excel. The difference in CT (dCT) values between the target and reference gene were calculated, using the formula $2^\wedge CT_{ref}/2^\wedge CT_{target}$, where $CT_{ref}$ = the CT value of the reference gene and $CT_{target}$ is the CT value of the target gene. The dCT values for each sample in a qPCR plate were normalized by dividing by the median of the entire plate. Data for each technical replicate for a given sample was normalized to the median of all dCT values for that patient to account for differences in baseline expression per patient (i.e. all technical replicates and all time-points analyzed). In some cases, CT values were not calculable.

For each patient and timepoint, a minimum of three normalized technical replicates was required, otherwise all values were omitted from further analysis, resulting in one fewer biological replicate for that timepoint. Outlier replicate data were identified and removed in GraphPad Prism using the Grubbs method (with $\alpha$ = 0.1%). The resulting mean relative abundance value of each remaining normalized technical replicate was considered a single biological replicate and was plotted using GraphPad Prism and included in the statistical analysis.

**Statistical Analysis of qPCR data.** Differences between active and placebo treatments were assessed per timepoint in GraphPad Prism (GraphPad Software, La Jolla California USA), using multiple t-tests, with adjustment for multiple comparisons using the Holm-Šídák method (alpha = 0.05; a consistent standard deviation was not assumed).

Microscopy Imaging: Cell images were taken with the Olympus Microscope IX71 and Olympus cellSens software.

#### SUPPLEMENTAL FIGURE CAPTIONS

**Supplemental Figure 1. (A)** SHP1705 structure. **(B)** Table of single dose SHP1705 pharmacokinetic parameters. **(C-E)** Gene expression levels of **(C)** core clock genes (*BMAL1/ARNTL*, *PER1*, *PER2*, *CRY1*, *RER-ERB*$\alpha$*/NR1D1*, *DPB*), **(D)** non-circadian, housekeeping genes (glyceraldehyde 3-phosphate dehydrogenase (*GAPDH*), Ribosomal protein lateral stalk subunit P0 (*RPLP0*), and Peptidylprolyl isomerase A (*PPIA*))**, (E)** blood transcriptome-based biomarker genes (Dishevelled associated activator of morphogenesis 2 (*DAAM2*), DNA damage inducible transcript 4 (*DDIT4*), Epoxide hydrolase 2 (*EPHX2*), Fatty acid amide hydrolase 2 (*FAAH2*), FOS-like 2, AP-1 transcription factor subunit (*FOSL2*), and Interleukin 6 (*IL6*)), in the presence and absence of SHP1705 administration (25, 75, or 500 mg, once-daily, morning administration). Data shown are mean ± SEM. N = 6 per treatment. Multiple t-tests (1 per time point), with Holm-Šídák adjustment for multiple comparisons. * $p < 0.05$

**Supplemental Figure 2. (A)** *CRY1* mRNA expression of non-tumor and GBM tissues from adult TCGA_GBM HG-U133A data plotted and analyzed with GlioVis. **(B)** Brightfield images of GSC 387, DGC 387, GSC 3565, DGC 3565, MGG 4, DGC 4, MGG 31, and DGC 31 at 10X magnification. **(C-G)** Cell viability analysis of NM cells (solid blue lines), GSCs (solid lines), or DGCs (dotted lines of matched GSC solid lines) **(C)** SHP1705, **(D)** KL001, **(E)** SHP656, or **(F)** SHP1703 treatment for 4 days (*n* = 3-4 biologically independent samples). **(G)** Heatmap of summarized $IC_{50}$ values following CRY activator treatment for 4 days in control NM cells, GSCs, and DGCs.

**Supplemental Figure 3.** Brightfield images of NM 290, GSC 3565, and MGG 31 at 10X magnification following 24hrs of DMSO or 1.25, 2.5, or 5 μM SHP1705 treatment.

**Supplemental Figure 4.** Average weight of mice injected with hGBM18 FMC MP1 and treated with for 1% CMC vehicle control, SHP1705 (10 mg/kg), and SHP1705 (30 mg/kg). Mice were weighed every 7 days (*n* = 10-11 mice/group).

**Supplemental Figure 5.** Cell viability analysis of NM cells, GSCs, or DGCs following **(A)** SR29065, **(B)** SR9009, or **(C)** SR9011 treatment for 4 days (*n* = 3-4 biologically independent samples). **(D)** Heatmap of summarized $IC_{50}$ values following REV-ERB agonist treatment for 4 days in control NM cells, GSCs, and DGCs.

### SUPPLEMENTAL FIGURES

**Supplemental Figure 1**

**A**

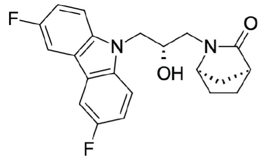

**B**

| Treatment | $C_{max}$ (ng/mL) | $C_{max}$ (hrs) | $AUC_{0\text{-}t}$ (ng*hrs/mL) | $T_{1/2}$ (hrs) |
|---|---|---|---|---|
| 500 mg SHP1705 Capsules Under Fasted Conditions | 24.1 ± 24.4 | 4 | 298 ± 203 | 24.7 ± 7.17 |

**C**

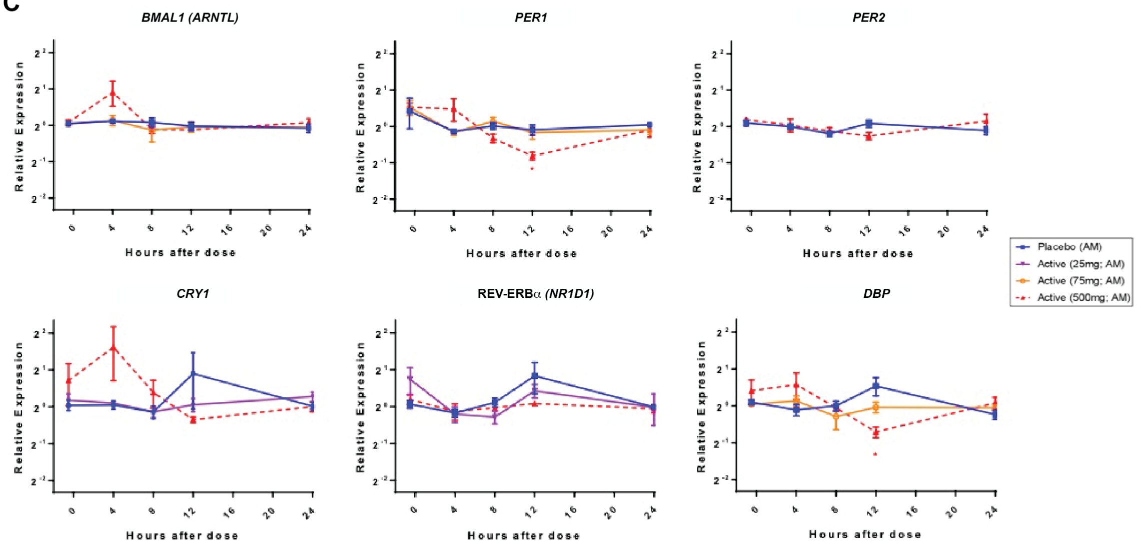

**D**

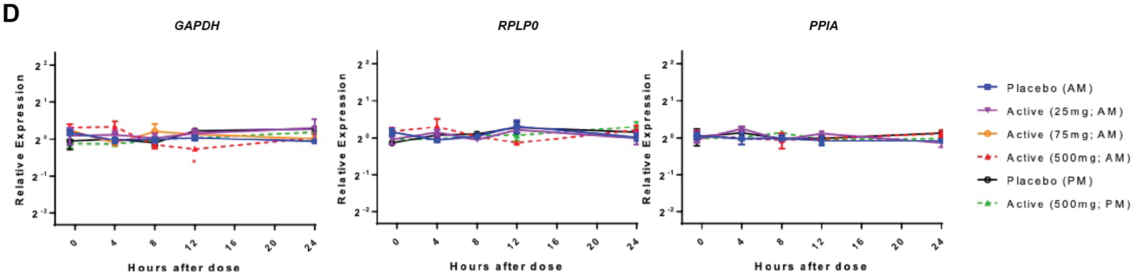

**E**

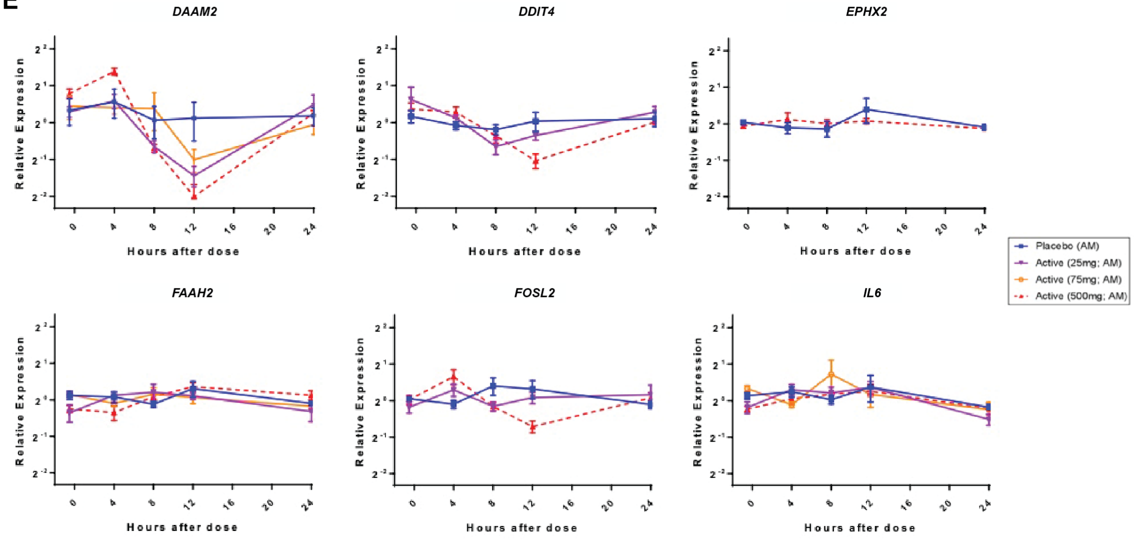

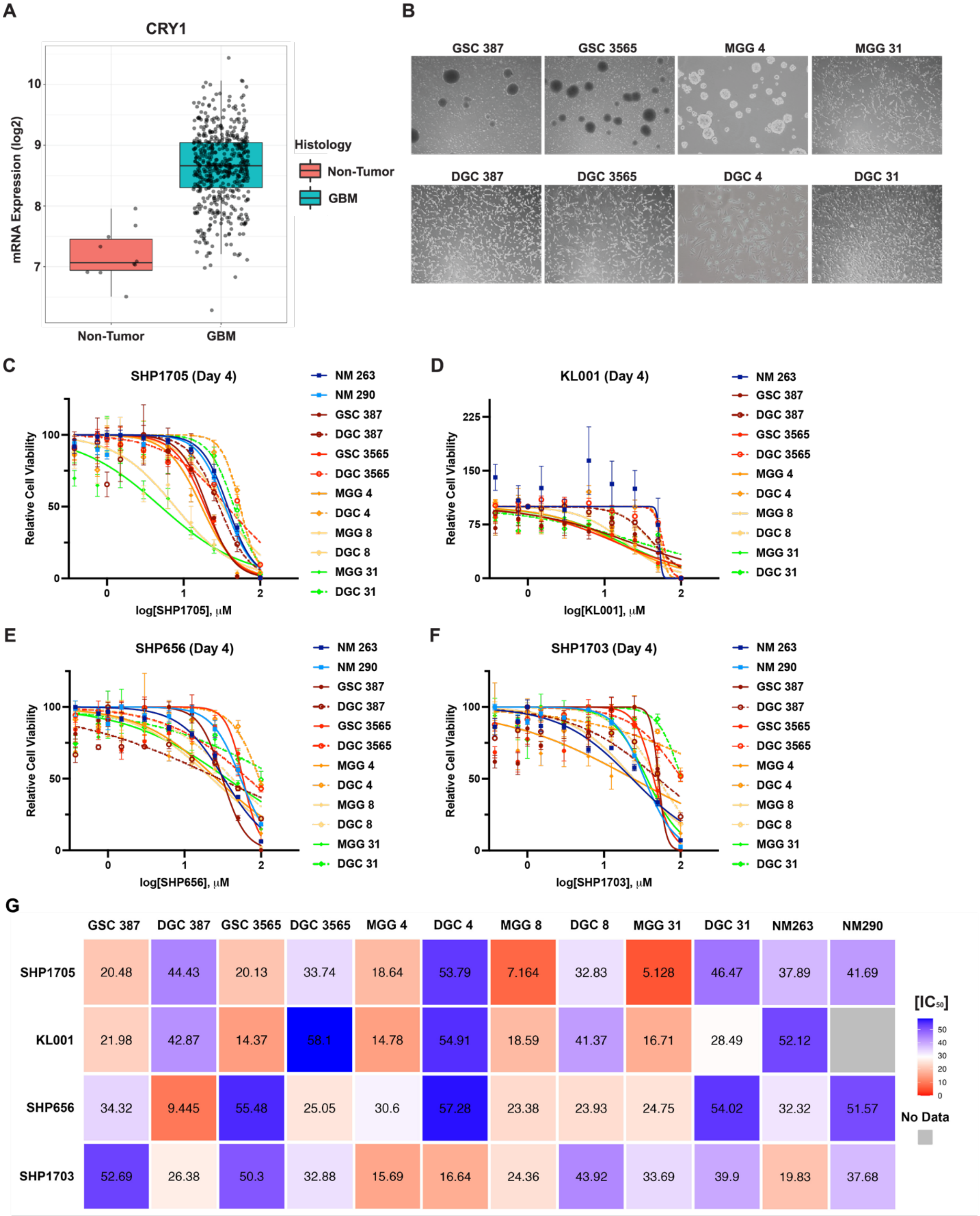

Supplemental Figure 2

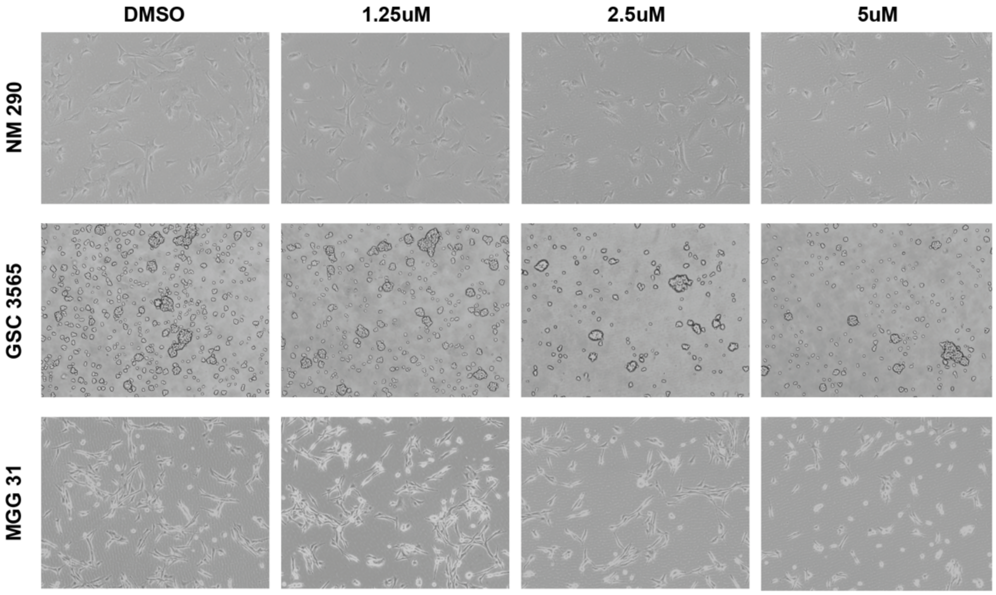

Supplemental Figure 3

Supplemental Figure 4

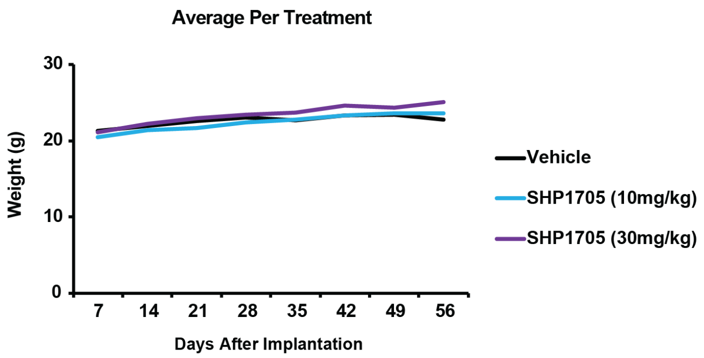

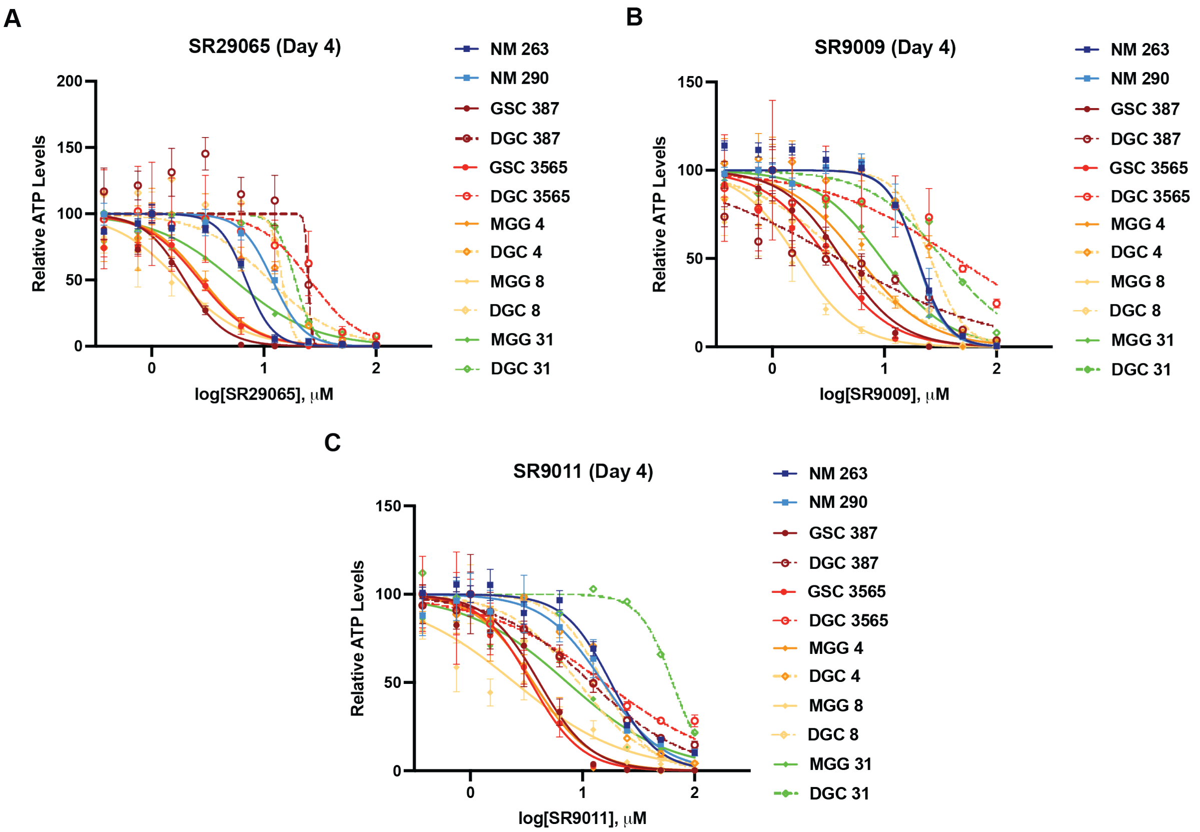

**D**

| | GSC 387 | DGC 387 | GSC 3565 | DGC 3565 | MGG 4 | DGC 4 | MGG 8 | DGC 8 | MGG 31 | DGC 31 | NM263 | NM290 |
|---|---|---|---|---|---|---|---|---|---|---|---|---|
| SR29065 | 1.958 | 20.7 | 2.658 | 25.62 | 2.822 | 11.15 | 1.737 | 14.58 | 5.388 | 18.92 | 6.921 | 11.92 |
| SR9009 | 3.888 | 14.67 | 2.999 | 18.2 | 5.341 | 20.17 | 1.667 | 4.272 | 8.358 | 21.99 | 16.62 | 18.28 |
| SR9011 | 3.372 | 7.146 | 2.994 | 4.633 | 3.608 | 14.64 | 2.216 | 8.635 | 5.77 | 27.76 | 13.66 | 11.71 |

$[IC_{50}]$ (color scale: 0, 10, 20)

**Supplemental Figure 5**

#### SUPPLEMENTAL TABLES

**Supplemental Table 1. Cell Lines Used in Cell Viability Assays**

| Cell Line | Malignancy/Status | TMZ Sensitivity |
|---|---|---|
| **NM 263** | Noncancerous | - |
| **NM 290** | Noncancerous | - |
| **GSC 387** | GBM, Stem Cell | N/A |
| **DGC 387** | GBM, Differentiated | N/A |
| **GSC 3565** | GBM, Stem Cell | N/A |
| **DGC 3565** | GBM, Differentiated | N/A |
| **MGG 4** | GBM, Stem Cell | Sensitive |
| **DGC 4** | GBM, Differentiated | Sensitive |
| **MGG 8** | GBM, Stem Cell | Sensitive |
| **DGC 8** | GBM, Differentiated | Sensitive |
| **MGG 31** | GBM, Stem Cell | Resistant |
| **DGC 31** | GBM, Differentiated | Resistant |
